# Highly efficient CRISPR gene editing in yeast enabled by double selection

**DOI:** 10.1101/262808

**Authors:** Philippe C Després, Alexandre K Dubé, Lou Nielly-Thibault, Nozomu Yachie, Christian R Landry

## Abstract

CRISPR-Cas9 loss of function (LOF) and base editing screens are powerful tools in genetics and genomics. Yeast is one of the main models in genetics and genomics, yet large-scale approaches remain to be developed in this species because of low mutagenesis rates without donor DNA. We developed a double selection strategy based on co-selection that increases LOF mutation rates, both for CRISPR-Cas9 and the Target-AID base editor. We constructed the pDYSCKO vector, which is amenable to high throughput double selection for both approaches. Using modeling, we show that this improvement provides the required increased in detection power to measure the fitness effects of thousands of mutations in typical yeast pooled screens. We also show that multiplex genome editing with Cas9 causes programmable chromosomal translocations at high frequency, suggesting that multiplex editing should be performed with caution and that base-editors could be preferable tools for LOF screens.

## Introduction

The CRISPR-Cas9 genome editing system has been engineered to create a comprehensive experimental toolkit (Jinek *et al.* 2012; Qi *et al.* 2013; Sander and Joung 2014). One of the principal applications of the system is for gene loss of function (LOF) via DNA repair errors within the coding sequence of a gene. The scalability of this technique allowed for the development of CRISPR-cas9 LOF experiments at the genome scale (Sanjana *et al.* 2014; Bassett *et al.* 2015; Sidik *et al.* 2016), greatly facilitating systems biology and genomics experiments in models in which they were difficult so far. Despite yeast being a usual frontrunner in technological developments for systems biology, low CRISPR-Cas9 LOF efficiency in yeast (Dicarlo *et al.* 2013) has so far made the application of this approach impractical. The power of the numerous resources already available for yeast would be greatly enhanced by the efficiency and cost-effectiveness of these new methods. For instance, it would allow for the generation of a fresh pool of LOF strains at the start of each experiment, minimizing secondary mutation accumulation, or even reconstituting a gene KO collection in a diversity of genetic backgrounds. This would be especially advantageous for strains that are used in industrial and biotechnological applications (Borodina and Nielsen 2014; Steensels *et al.* 2014) and in the context of natural strains for the study of evolution (Marsit *et al.* 2017). Newly developed base editors based on nCAs9 and dCas9 fusions, such as Target-AID (Nishida *et al.* 2016), which induces C to G and C to T changes (with rare C to A), would also make important additions to the yeast systems biology toolkit if they could be used with high mutagenesis efficiency.

One of the crucial determinant of large-scale LOF screens is mutagenesis efficiency. Accordingly, different approaches have been used to optimize mutagenesis rate. One of them is co-selection, which relies on the selection of cells based on a marker that has mutated alongside the target loci based on a positively or negatively selected marker. This was recently shown to enhance edition rates in human cell lines (Agudelo *et al.* 2017) and drosophila (Kane *et al.* 2017), but has not been applied to yeast yet. Because the base efficiency of CRISPR-Cas9 KO in these model organisms is higher than in yeast by several orders of magnitudes (Dicarlo *et al.* 2013), the gain of efficiency brought by co-selection in yeast would be even greater than what it is in these other models. Co-selection has also be shown to enhance base-editing efficiency in human cell lines (Billon *et al.* 2017).

We developed a double selection system for yeast to exploit this principle. We present a high efficiency co-selection approach to easily produce LOF mutants in yeast with both CRISPR-Cas9 and Target-AID using vectors amenable to genome-wide screens. It is intuitive that a higher efficiency would allow for more efficient screening as well as the detection of more subtle effects on fitness. However, it is not clear by how much we need to improve editing efficiency to be able to perform experiments in yeast using CRISPR-Cas9 LOF or base editing so that they would compare in terms of power with experiments currently performed with the yeast deletion collection. We therefore developed a statistical model to quantitatively assess the improvement brought by double selection on CRISPR-LOF screens and use it to show that it is highly significant, particularly for fitness effects in the range in which most of gene deletion effects are observed across a range of growth conditions.

## Results and discussion

The double selection system uses the pDYSCKO (Double Yeast Selection CRISPR-KO) plasmid, which encodes for two guide RNAs (gRNA), one targeting the gene of interest (gYFG) and the other the negative selection market *CAN1* (Figure 1). Another vector is used for galactose induction of the effector enzyme, which can be different variants of Cas9. LOF of *CAN1* confers resistance to the antibiotic canavanine, which means that canavanine media can be used to enrich the cell population for mutant cells.

**Figure 1.**
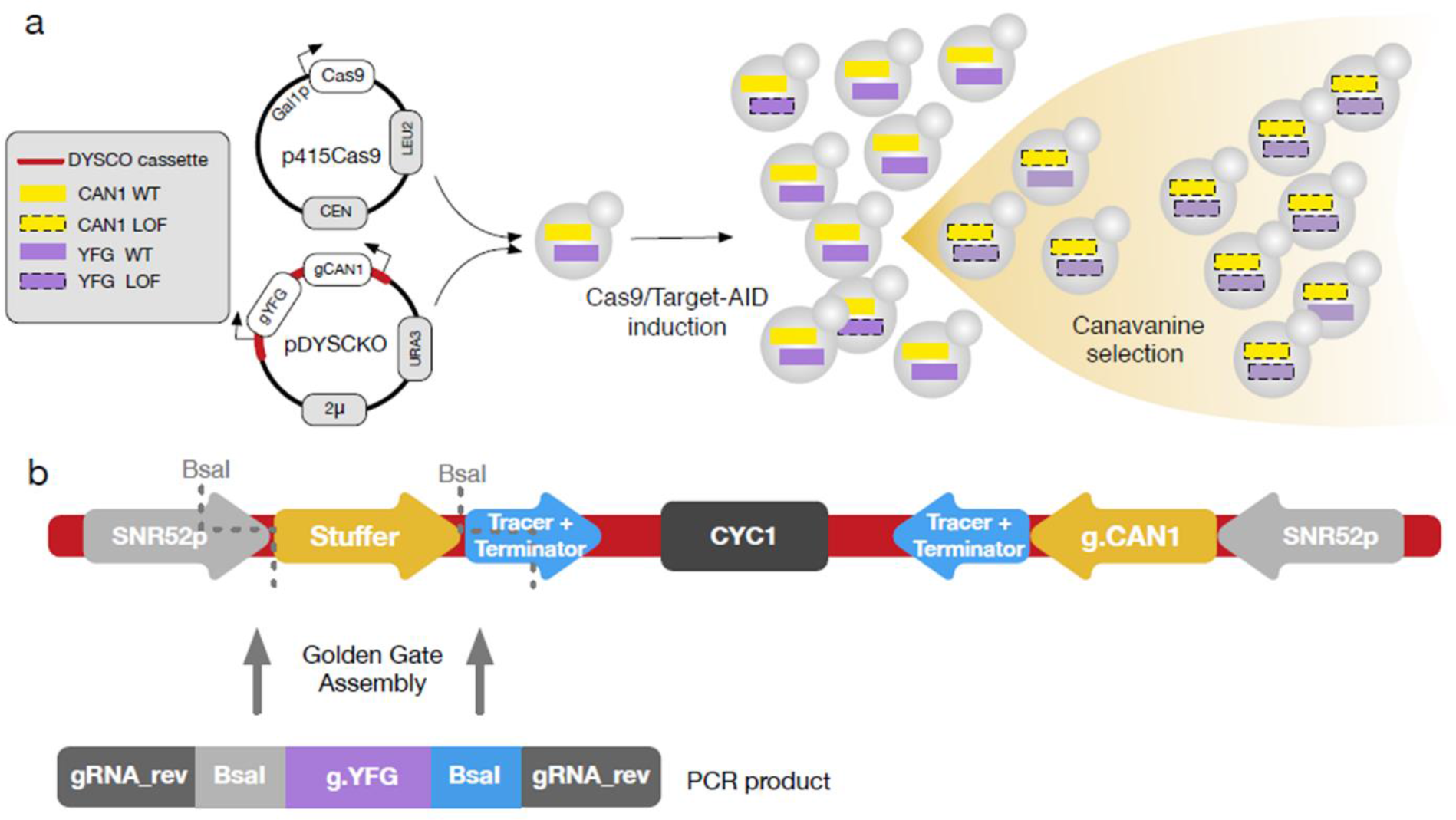
The pDYSCKO-CAN1 plasmid for double selection LOF. a) Overview of co-selection using the pDYSCKO and the p415Cas9 plasmid, expressing guide RNAs and Cas9 respectively. Only a fraction of cells are successfully edited during Cas9 induction (galactose media), but cells with a LOF at one locus have a greater chance of bearing LOFs at both loci. Canavanine selection allows for enrichment for these cells. b) Overall structure of the DYSCKO cassette, which is on a plasmid also containing standard selection markers (AmpR and URA3). The two symmetrical gRNA expression units use the same promoters and terminators (from SNR52 and SUP4 respectively), insuring their co-expression. The stuffer is a short sequence containing two restriction sites for BsaI that does not match any sequence in the yeast genome. Custom gRNA insertion in the vector is performed through Golden Gate assembly (Engler *et al.* 2008) using a short dsDNA fragment containing the gRNA target the sequence of interest (g.YFG).

We estimated mutation rates using a gRNA targeting the *ADE1* gene, which leads to a red colony phenotype when inactivated, allowing for direct LOF rate estimates. We compared conditions with and without double selection. The baseline KO efficiency without double selection confirms the initial report (Dicarlo *et al.* 2013) of less than 0.1%. The *ADE1* LOF rate is far greater for both CRISPR-Cas9 and Target-AID (Figure 2) when co-selected using the *CAN1* marker. The median mutation rate for CRISPR- Cas9 shows a 1200-fold increase (median mutation rate: 62%). Target-AID is also improved, with median fold increase of 3 (median mutation rate: 61%). The improvement is relatively modest but the basal rate is already relatively high, which brings the LOF rate well above 50%. In addition, we note that the actual edition rate is most likely underestimated. Target-AID performs C to T and C to G edition (with rare C to A) (Nishida *et al.* 2016) and we find that white colonies after recovery systematically show C to T mutations (Figure 2b), making them silently changed with respect to the phenotype. The 60% LOF rate could therefore correspond to about 100% editing rate. We also measured the LOF efficiency using pDYSCKO in diploid cells. While Target-AID mutagenesis proved efficient (median mutation rate 45%, Figure 2a), Cas9 mediated LOF was highly toxic and the efficiency was lower than in haploids by several orders of magnitude. LOF mutant construction in diploid cells should therefore use the Target-AID system.

**Figure 2.**
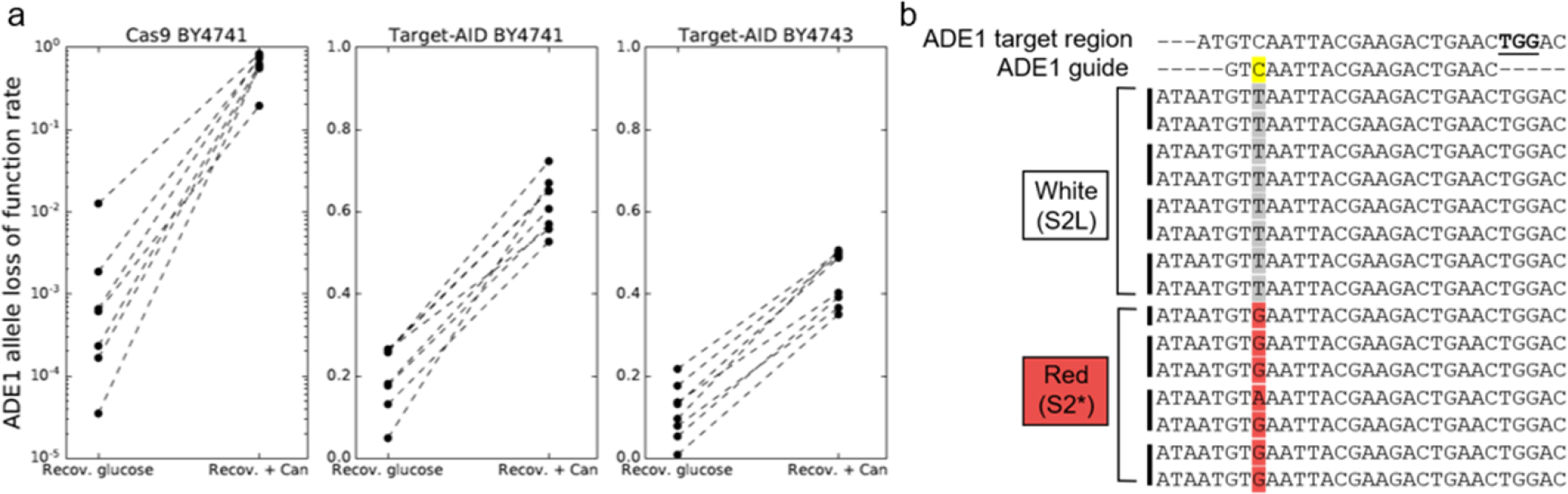
Double selection increases gene editing efficiency. a) The same cell cultures were placed in different recovery conditions after induction of Cas9 or Target-AID expression. Recovery in glucose is neutral with respect with the *CAN1* mutant genotype. Recovery in canavanine selects for cells that have LOF mutations at the *CAN1* locus. Selection on the *CAN1* marker increases the representation of ADE1 LOF colonies. Cells with non-functional *ADE1* alleles accumulate a red metabolic intermediate that gives a characteristic color to colonies allowing for easy identification of mutants, while mutants without LOF mutations in *ADE1* have a wild-type phenotype. The left panel shows standard Cas9 KO in haploid cells, the middle panel Target-AID base edition in haploid cells and the right panel, Target-AID edition in diploid cells. Plating was performed after a 16-hour recovery period. b) Mutations induced by Target-AID in white and red colonies from four independent experiments. The base editing site is shown in yellow in the guide sequence. The changes induced correspond to those expected, with similar proportions of C-T and C-G and rare C-A (Nishida *et al.* 2016). All white colonies have been edited as well, showing that the LOF only occurs with the C to G mutations and thus cannot reach a rate of 100% in these specific assays.

In a typical LOF screen, pools of gRNA are cloned and transformed in a population of cells. The fate of individual mutants is followed through competition assays in which gRNA sequences serve as barcode to estimate the abundance of each genotype individually by deep sequencing, which is analogous to yeast Bar-seq approaches with the yeast deletion collection (Robinson *et al.* 2014). This approach relies on high mutation rate because a population of cells bearing a specific gRNA could be a mixture of mutants and WTs, which cannot be distinguished from sequencing the gRNA barcodes alone. We can therefore use existing data on Bar-seq-based fitness measurements to examine how the improved efficiency of gene KO and editing afforded by the double selection can increase the detection power of small fitness effects.

Our simulations show that the increased efficiency allows for more sensitive detection of LOF effects (Figure 3), particularly in the critical zone encompassing fitness effects between 0% and 25%, which are the most frequent effects for the deletion of non-essential genes in yeast. Our double selection strategy therefore makes genome-wide screening powerful enough to meet the high standards established by the yeast deletion collection. The model also predicts that the efficiency of Target-AID editing in diploid cells would be high enough to consider large-scale screening applications. For example, at a mutagenesis rate of 0.2, a LOF with a selection coefficient of 0.05 has a 25% chance of being detected, while a rate of 0.6 instead allows it to be detected in over 99% of cases.

**Figure 3.**
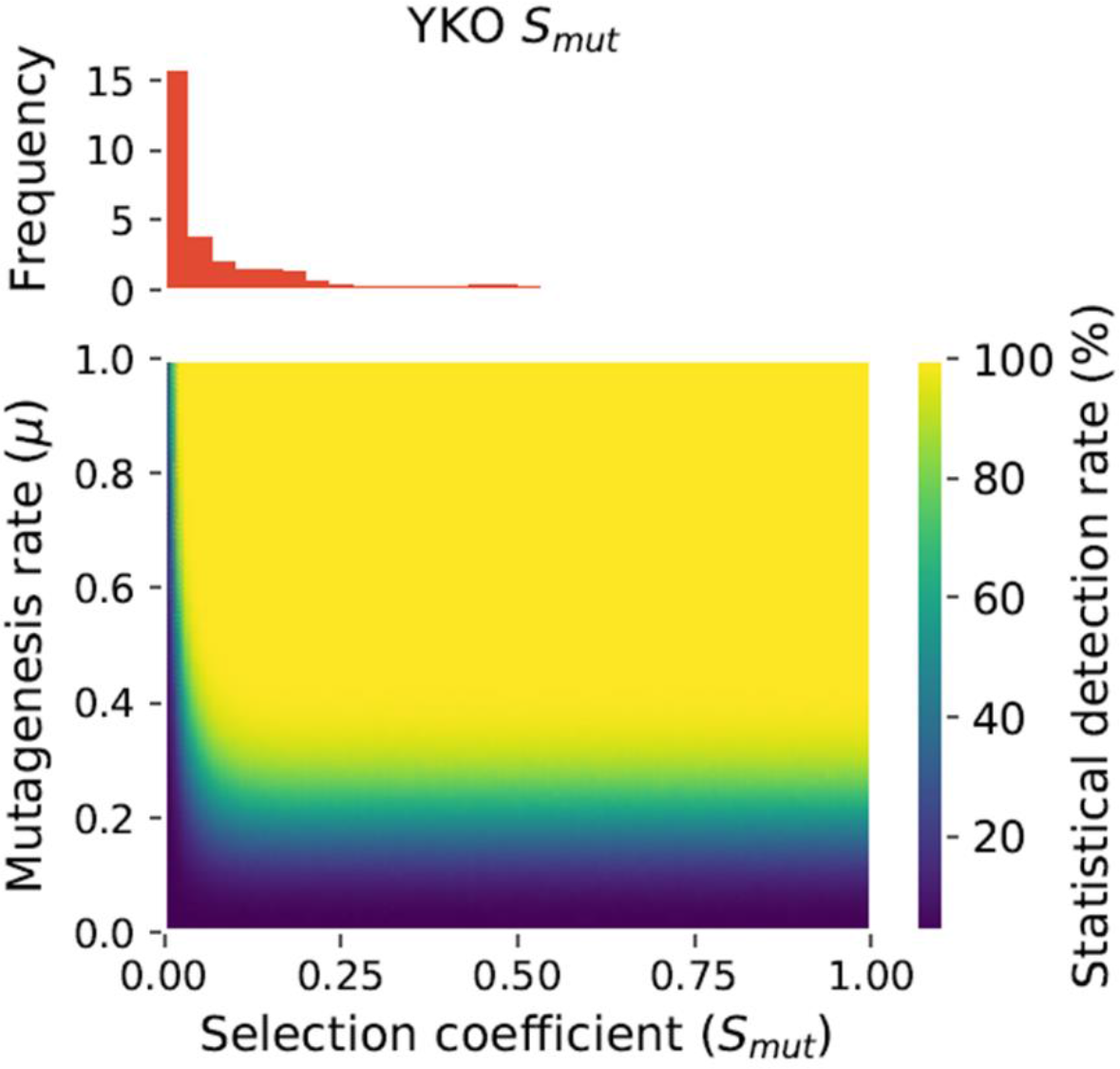
Improvement in mutagenesis rates enhance the power of CRISPR-LOF screens. The rate of detection of fitness defects is shown as a function of mutagenesis rate and the selection coefficient of the mutation. Improving the mutation rate above 0.3 dramatically increases the ability to detect growth defects, a requirement in many experiments including genome-wide CRISPR LOF barcode sequencing. The increase is particularly noticeable for selection coefficient associated to the majority of gene KO (YKO) mutants in various conditions (Qian *et al.* 2012), as shown in the panel above.

Our approach relies on the *CAN1* gene that can be used for negative selection against non-mutated cells. This marker has been used in many large-scale and systematic experiments, for instance synthetic gene arrays (SGA) (Tong *et al.* 2001), which means that the strains constructed are compatible in terms of genotype with other approaches routinely used by the yeast community. If needed, other negatively selectable markers could be exploited, for instance *LYP1, FCY1* and *URA3.* Similarly, the plasmid selection markers can easily be changed to antibiotic resistance cassettes for G418 (Wach *et al.* 1994), Nourseothricin or Hygromycin B (Goldstein and McCusker 1999), making it a polyvalent tool for high throughput gene disruption in non-standard laboratory strains.

Our experiment also revealed that multiplexed gene CRISPR-Cas9 LOF in yeast has undesirable side effects. We recurrently observed colonies exhibiting an intermediate phenotype for *ADE1* LOF (median rate: 18%), with a color pattern slightly different from the standard red *ade1* phenotype. We found that these cells harbored a chromosomal rearrangement that results into a *CAN1-ADE1* fusion. We first identified these fusions by PCR and Sanger sequencing and confirmed that they affect chromosome size by PFGE (Figure 4a, 4b). We find that changing the DYSCKO target site from ADE1 to VPS35 produced a different fusion but at a similar rate (Figure 4c), showing that these fusions are likely programmable and generalizable.

**Figure 4.**
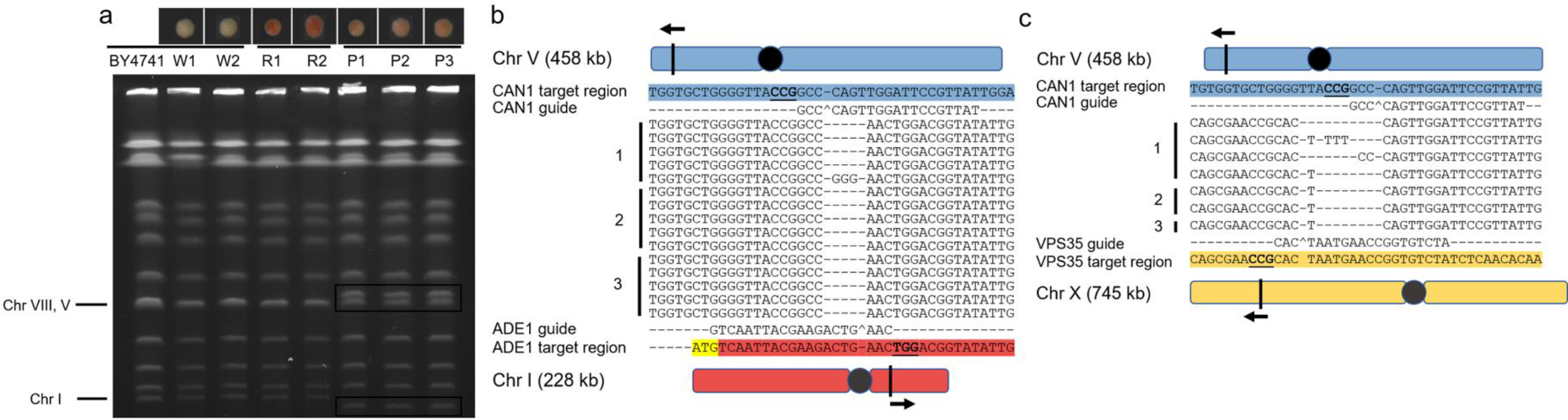
Multiplex CRISPR-Cas9 causes chromosomal rearrangements in yeast. a) Chromosomal migration patterns of the different canavanine resistant mutants generated using CRISPR-Cas9. Mutants without LOF mutations in *ADE1* have the same phenotype as the wild-type strain. Colonies with a distinct pink color show chromosomal fusions. *CAN1* only LOF (W1 and W2) and red *CAN1, ADE1* LOF mutants show a chromosomal migration pattern that is comparable to the wild-type. Pink colonies from independent mutagenesis experiments show altered length for chromosomes I and chromosomes V, consistent with a reciprocal translocation centered on the predicted Cas9 cut sites. b) Breakpoint sequencing of 5 pink colonies from 3 independent mutagenesis experiments confirms PFGE patterns. c) CRISPR-Cas9 mutagenesis with a gRNA targeting *VPS35.* Colonies with fusion alleles were identified by PCR. Breakpoint sequencing of 7 chromosomal translocations from three independent experiments confirmed the translocations. Fusion occurrences per replicate were 4 (4) /15, 3 (2) /16, and 1 (1) /16, median rate 18.75%. The number in parenthesis represents the number of detected breakpoints that were sequenced

The observation that multiplex genome editing causes chromosomal rearrangements at relatively high frequency opens new opportunities for the use of genome engineering in yeast and likely other organisms as well. It is however also worrisome because many recent multiplex mutagenesis workflow rely on PCR based deep sequencing of the target allele to measure success rates (i.e. CRISPR-Seq (Tothova *et al.* 2017)) of genome editing at multiple loci. This method cannot detect chromosomal rearrangements, and as such the events may have gone unnoticed in many experiments so far. It would have been in our case if we did not have access to visible phenotypes. It is unknown if the high occurrence rate is specific to yeast, but efforts should be made to assess it in other models. As we have yet to observe any case of rearrangements when using Target-AID, base editors might be safer choices in most cases when attempting multiplex LOF mutagenesis. One possible tradeoff with these editors is that some of the induced changes may lead to silent substitutions, such as the ones we observed here, which effectively limit the effective LOF rate. This tradeoff can be alleviated by a careful design of gRNA in order to maximize the effects of the induced base substitution. The development of other base editors, for instance from A-T to G-C (Gaudelli *et al.* 2017), offer new possibilities in terms of producing loss of function mutations without DNA cleavage, diminishing even more the need to use systems that may lead to chromosomal fusions.

## Methods

The pDYSCKO vector was built from p426-SNR52p-gRNA.CAN1.Y-SUP4t (Addgene # 43803). Directed mutagenesis was used to create silent mutations in the AmpR gene and the *URA3* gene to remove BsaI restriction sites using the oligonucleotides mut_AmpR_For/mut_AmpR_Rev and mut_URA3_For/mut_URA3_Rev (all oligonucleotides used in this study are presented in Table S1). The DYSCKO cassette was synthetized as a gBLOCK (Integrated DNA Technologies, Coralville, USA) and Gibson cloned into the modified backbone amplified with Backbone_For and Backbone_Rev. The g.ADE1 ssDNA oligonucleotide containing the gRNA targeting the *ADE1* gene (or *VPS35*) was amplified by PCR using the gRNA_for and gRNA rev oligonucleotides. This guide was previously used by Nishida et al (Nishida *et al.* 2016) to create LOFs in *ADE1* using Target-AID and is also well suited to CRISPR-Cas9 gene disruption. Multiple amplification reactions were pooled and purified using the EZ-10 Column PCR Products Purification Kit (Biobasic, Markham, Canada). Fragment concentration was estimated using a NanoDrop (Thermofisher, Waltham, USA). The insert was cloned into pDYSCKO using the Golden Gate Assembly Mix (New England Biolabs, Ipswich, USA) with the following parameters: 1 ul pDYSCKO vector (50 ng/ul), 1 ul insert (0.5 ng/ul) in the standard manufacturer 20 ul assembly reaction conditions, which was incubated for 1 hour at 37°C followed by 5 minutes at 55°C. The plasmids were transformed in bacteria and positive clones were confirmed by Sanger sequencing (CHUL sequencing platform, Québec, Canada) using the DYSCKO_rev oligonucleotide.

All experiments were performed in the haploid S. *cerevisiae* BY4741 strain (MATa his3Δ1 leu2Δ0 met15Δ0 ura3Δ0) or diploid BY4743 (MATa/α his3Δ1/his3Δ1 leu2Δ0/leu2Δ0 LYS2/lys2Δ0 met15Δ0/MET15 ura3Δ0/ura3Δ0) from (Baker Brachmann *et al.* 1998). Plasmid p415-Cas9 was obtained from Addgene (#43804). The nCas9-Target-AID plasmid was a generous gift from Dr Keiji Nishida, Kobe University, Kobe, Japan.

Media recipes are detailed in Table S2. All yeast growth occurred at 30°C and with shaking in the case of liquid cultures._Competent cells and transformations were performed using standard protocols (Amberg *et al.* 2005), with a two-hour recovery period. Cells were plated on SC-UL and allowed to grow for 48 hours at 30°C before the start of mutagenesis experiments.

Multiple transformation colonies were used during mutagenesis to inoculate a 3 ml SC-UL + 2% glucose culture in which cells were grown for 24 hours. Enough cells to inoculate at 1 OD600 a 3ml culture were harvested by centrifugation and placed in SC-UL+5% glycerol for 24 hours. Cas9 or Target-AID expression was then induced by switching the media to 3 ml SC-UL+ 5% galactose for a 12-hour period. Cells were diluted to OD of 0.1 in 3 ml SC-ULR +2% glucose + Canavanine (50 µg/ml) for a 16- hour double selection or 3 ml SC-UL +2% glucose at the same OD as the control condition.

Mutation rates were assessed by plating cells on SC-ULR + Can and SC-ULR after galactose induction and after recovery with or without double selection at appropriate dilutions. Mutation rate was assessed by calculating the ratio of red (or pink) colonies over total number of colonies. The *ADE1* and *CAN1* alleles of 8 white and 8 red colonies were sequenced to confirm mutations either through Cas9 or Target-AID with CAN1_for, CAN1_rev and ADE1_for, ADE1_rev. The CAN1_For and ADE1_Rev oligonucleotides was used to amplify the *CAN1-ADE1* breakpoint and the resulting amplicon was Sanger sequenced (CHUL sequencing platform, Québec, Canada).

Chromosomal rearrangements were confirmed by PFGE (Maringele and Lydall 2006). Plugs were prepared using cells from overnight cultures in YPD for BY4741 and SC- ULR + canavanine for mutant strains. Migration time was 27 hours, with a switch time of 60 seconds and no ramping.

Fusions with the *VPS35* gene were engineered using the same approach but with pDYSCKO-*VPS35.* Fusions were detected by PCR using the VPS35-A and CAN1_rev oligonucleotides, and the resulting amplicons were Sanger sequenced (CHUL sequencing platform, Québec, Canada)._Sequence were aligned using the MEGA 7 software (Kumar *et al.* 2016).

In CRISPR-LOF screens, populations of cells are transformed with pools of vectors bearing different gRNAs which are amplified by PCR and used as barcode in a barcode-sequencing competition experiment (Bar-seq) (Shalem *et al.* 2014; Robinson *et al.* 2014), the gRNA being associated to genes by sequence identity. We modeled this process by adapting haploid selection models, with the added challenge that not all cells with a specific guide will bear a LOF at the target locus because mutation rate is not 100%.

Let *n_WT_* and *n_BC_* be respectively the abundance of a control guide and the abundance of a guide targeting a gene which when inactivated has a selection coefficient *S_mut_.* This mutation is present in a proportion *μ*, which represents the mutation rate after recovery and thus the frequency at the beginning of the competition. After a time *t,* the abundance ratio of these two barcodes is described by

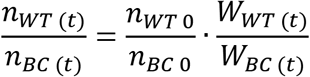

Were *W_WT(t)_* is the absolute fitness of the control barcode population at time *t* and *W_BC(t)_* the absolute fitness of the barcode associated with the mutation. Because the rate of mutation is not 100% for the strains harboring the barcode (*µ* < 1), a certain proportion of cells (1 − *μ*) will have the same fitness as cells bearing the control barcode so that:

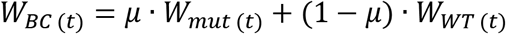

and

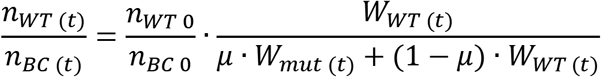

By dividing both the numerator and the denominator of the rightmost factor by *W_WT(t)_*, we obtain

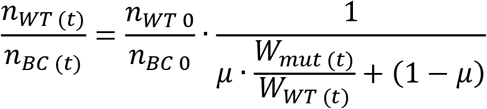

And then to

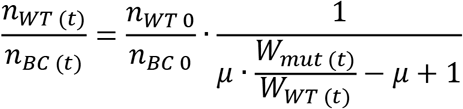

Because we assume that the growth rates of the mutants and wild-type are constant over time:

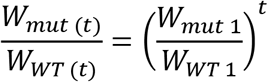

Given that the selection coefficient *S_mut_* is 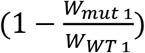 (Hartl 2000) we have

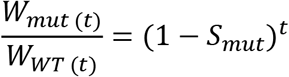

And therefore

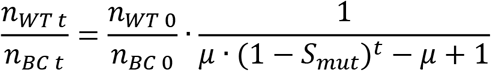

Which is simplified to

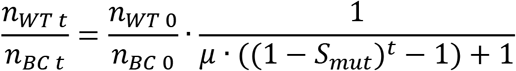

If we assume that the control barcode and the mutant are present in the same proportion at the start of the experiment, then 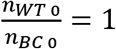 and

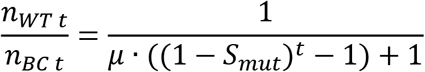

This gives us a theoretical population ratio at time *t* for the control barcode over the barcode for a mutant, as a function of the mutagenesis rate and the selection coefficient over a generation. Because we observe these ratios by sequencing the barcode pool at a given depth, the number of reads for each barcode will be influenced by stochastic sampling. If we have *k* guides in the experiment, and sequence at an estimated depth of *d* reads on average per guide, and we assume all barcodes have equivalent abundance at the start, then the number of reads *R* sequenced for a barcode at *t* = 0 follows

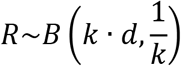

Because of our previous assumptions, this distribution is valid at *t* = 0 for all barcodes regardless of μ or *S_mut_*. For other values of *t*, if we assume that the distribution remains equivalent for the controls barcodes, then *R* for a LOF barcoded strain will follow a binomial distribution influenced by the abundance change caused by the LOF of the target allele:

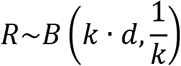

Where

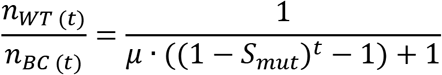

We can therefore obtain simulated sequencing read number distributions as a function of *μ*, *S_mut_*, *t*, *k*, and *d*. To assess whether a difference in read count for a barcode is significant, we can test for equality of proportion between a control barcode and the barcode of interest using the chi-square test, with the following table:

|  |  |
| --- | --- |
| $R_{MutBC\ 0}$ | $R_{MutBC(t)}$ |
| $R_{wt\ 0}$ | $R_{wt\ (t)}$ |

Such experiments include multiple control gRNA that can be pooled in the same category. Most of the time, there are also multiple guides targeting the gene. However, because Target-AID can be used to create specific point mutations that may not all be equivalent to LOF and may not have the same efficiency, it is useful to consider all guides individually. If we set the number of guide per target loci to *g*, the number of target loci to *t*, and the number of control *c*, then the total sequencing depth for a set *d* changes because:

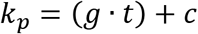

Because we expect all controls to be equivalent, then we can assume that

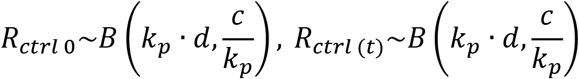

And for a single target

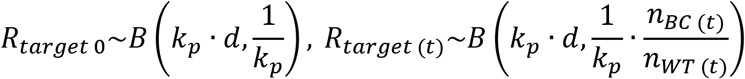

We used this model to simulate a CRISPR-LOF screen with parameters analogous to a Bar-seq screen in yeast. We draw simultaneously a sample from each distribution and test the values for significance over 10 000 iterations. This was done for 100 increments of *μ* from 0 to 1 and 100 increments of *S_mut_* from 0 to 1.

The yeast genome contains ∼6,000 annotated ORFs, of which 1,156 are essential and not useful for a standard LOF screen. Control loci could include intron sequences, pseudogenes or predicted putative non-functional peptides, or often used controls like the HO nuclease. Therefore, to simulate a CRISPR-LOF screen in yeast, the following parameters were used *t* = 4,800 (number of LOF target loci), *c* = 800 (number of control guides), *g* = 8 (number of guides per target loci), *d* = 200 (expected average coverage per guide) and *t* = 26 (number of generations). The CRISPR-Cas9 LOF simulation was performed using a script written in python 2.7 using the Ipython/Jupyter interactive environment (v4.1.1) (Perez and Granger 2007), Numpy (v1.13.3) (van der Walt *et al.* 2011), Pandas (v0.20.3) (McKinney 2010), Matplotlib (v2.1.0) (Hunter 2007) and SciPy (v0.13.3) (Oliphant 2007). Fitness effects of gene deletions in Bar-seq experiments were retrieved from Qian et al (Qian *et al.* 2012) and pooled across experiments.

## Availability

The pDYSCKO vector will be deposited on Addgene (http://www.addgene.org/), as well as pDYSCKO-*ADE1*. The Python 2.7 Packages used are available through in the Python Package Index (https://pypi.python.org/pypi/pip) or in Conda (https://conda.io/docs/).

## Supplementary material

Oligonucleotides used in this study are presented in table S1. Media recipes are presented in table S2.

## Acknowledgements

The authors thank G. Charron for help with the PFGE protocol, and M. Hénault and D. Yamamoto Evans for insightful discussions.

## Author Information

Corresponding Author: Christian Landry. Room 3106, Pavillon Charles-Eugène- Marchand, 1030, Avenue de la Médecine, Université Laval, Québec (Québec) G1V 0A6, Canada. Tel. 1-418-656-3954, Fax 1-418-656-7176

## Contributions

P.C.D and A.K.D performed the experiments. P.C.D and A.K.D designed research with support from C.R.L. and N.Y. P.C.D. developed the model with L.N.T. P.C.D analyzed the data and wrote the manuscript with contributions from all authors.

## Funding

This work was supported by the Canadian Institutes of Health Research [299432, 324265 to C.R.L. 364920, 384483 and Frederick Banting and Charles Best graduate scholarship to P.C.D.], the Natural Sciences and Engineering Research Council [Alexander Graham Bell graduate fellowship to L.N.T.], Laval University [André Darveau Fellowship to P.C.D.] and the Japan Society for the Promotion of Science [S15734 and S17161 to C.R.L. and N.Y.].

## Conflict of interest

The authors declare no competing financial interests.

## References

Agudelo D., Duringer A., Bozoyan L., Huard C. C., Carter S., et al., 2017 Marker-free coselection for CRISPR-driven genome editing in human cells. Nat. Methods 14: 615–620.

Amberg D. C., Burke D. J., Strathern J. N., 2005 Methods in Yeast Genetics: A Cold Spring Harbor Laboratory Course Manual, 2005 Edition.

Baker Brachmann C., Davies A., Cost G. J., Caputo E., Li J., et al., 1998 Designer deletion strains derived fromSaccharomyces cerevisiae S288C: A useful set of strains and plasmids for PCR-mediated gene disruption and other applications. Yeast 14: 115–132.

Bassett A. R., Kong L., Liu J.-L., 2015 A genome-wide CRISPR library for high-throughput genetic screening in Drosophila cells. J. Genet. Genomics 42: 301–9.

Billon P., Bryant E. E., Joseph S. A., Nambiar T. S., Hayward S. B., et al., 2017 CRISPR-Mediated Base Editing Enables Efficient Disruption of Eukaryotic Genes through Induction of STOP Codons. Mol. Cell.

Borodina I., Nielsen J., 2014 Advances in metabolic engineering of yeast Saccharomyces cerevisiae for production of chemicals. Biotechnol. J. 9: 609–620.

Dicarlo J. E., Norville J. E., Mali P., Rios X., Aach J., et al., 2013 Genome engineering in Saccharomyces cerevisiae using CRISPR-Cas systems. Nucleic Acids Res. 41: 4336–4343.

Engler C., Kandzia R., Marillonnet S., 2008 A one pot, one step, precision cloning method with high throughput capability. PLoS One 3.

Gaudelli N. M., Komor A. C., Rees H. A., Packer M. S., Badran A. H., et al., 2017 Programmable base editing of A•T to G•C in genomic DNA without DNA cleavage. Nature 551: 464–471.

Goldstein A. L., McCusker J. H., 1999 Three new dominant drug resistance cassettes for gene disruption inSaccharomyces cerevisiae. Yeast 15: 1541–1553.

Hartl D. L., 2000 A Primer of Population Genetics. Sunderland, Mássachussets. U.S.A Sinauer Assoc. Inc.: 26–31.

Hunter J. D., 2007 Matplotlib: A 2D Graphics Environment. Comput. Sci. Eng. 9: 90–95.

Jinek M., Chylinski K., Fonfara I., Hauer M., Doudna J. A., et al., 2012 A Programmable Dual-RNA-Guided DNA Endonuclease in Adaptive Bacterial Immunity. Science (80-.). 337: 816–821.

Kane N. S., Vora M., Varre K. J., Padgett R. W., 2017 Efficient Screening of CRISPR/Cas9-Induced Events in Drosophila Using a Co-CRISPR Strategy. G3 (Bethesda). 7: 87–93.

Kumar S., Stecher G., Tamura K., 2016 MEGA7: Molecular Evolutionary Genetics Analysis Version 7.0 for Bigger Datasets. Mol. Biol. Evol. 33: 1870–1874.

Maringele L., Lydall D., 2006 Pulsed-field gel electrophoresis of budding yeast chromosomes. Methods Mol. Biol. 313: 65–73.

Marsit S., Leducq J. B., Durand É., Marchant A., Filteau M., et al., 2017 Evolutionary biology through the lens of budding yeast comparative genomics. Nat. Rev. Genet. 18: 581–598.

McKinney W., 2010 Data Structures for Statistical Computing in Python. : 51–56.

Nishida K., Arazoe T., Yachie N., Banno S., Kakimoto M., et al., 2016 Targeted nucleotide editing using hybrid prokaryotic and vertebrate adaptive immune systems. Science 102: 553–563.

Oliphant T. E., 2007 SciPy: Open source scientific tools for Python. Comput. Sci. Eng. 9: 10–20.

Perez F., Granger B. E., 2007 IPython: A System for Interactive Scientific Computing. Comput. Sci. Eng. 9: 21–29.

Qi L. S., Larson M. H., Gilbert L. A., Doudna J. A., Weissman J. S., et al., 2013 Repurposing CRISPR as an RNA-guided platform for sequence-specific control of gene expression. Cell 152: 1173–83.

Qian W., Ma D., Xiao C., Wang Z., Zhang J., 2012 The Genomic Landscape and Evolutionary Resolution of Antagonistic Pleiotropy in Yeast. Cell Rep. 2: 1399–1410.

Robinson D. G., Chen W., Storey J. D., Gresham D., 2014 Design and analysis of Bar-seq experiments. G3 (Bethesda). 4: 11–8.

Sander J. D., Joung J. K., 2014 CRISPR-Cas systems for editing, regulating and targeting genomes. Nat. Biotechnol. 32: 347–55.

Sanjana N. E., Shalem O., Zhang F., 2014 Improved vectors and genome-wide libraries for CRISPR screening. Nat. Methods 11: 783–784.

Shalem O., Sanjana N. E., Hartenian E., Shi X., Scott D. A., et al., 2014 Genome-scale CRISPR-Cas9 knockout screening in human cells. Science (80-.). 343: 84–87.

Sidik S. M., Huet D., Ganesan S. M., Huynh M.-H., Wang T., et al., 2016 A Genome-wide CRISPR Screen in Toxoplasma Identifies Essential Apicomplexan Genes. Cell 166: 1423–1435.e12.

Steensels J., Snoek T., Meersman E., Nicolino M. P., Voordeckers K., et al., 2014 Improving industrial yeast strains: Exploiting natural and artificial diversity. FEMS Microbiol. Rev. 38: 947–995.

Tong A. H., Evangelista M., Parsons A. B., Xu H., Bader G. D., et al., 2001 Systematic genetic analysis with ordered arrays of yeast deletion mutants. Science 294: 2364–8.

Tothova Z., Krill-Burger J. M., Popova K. D., Landers C. C., Sievers Q. L., et al., 2017 Multiplex CRISPR/Cas9-Based Genome Editing in Human Hematopoietic Stem Cells Models Clonal Hematopoiesis and Myeloid Neoplasia. Cell Stem Cell 21: 547–555.e8.

Wach A., Brachat A., Pöhlmann R., Philippsen P., 1994 New heterologous modules for classical or PCR-based gene disruptions in Saccharomyces cerevisiae. Yeast 10: 1793–808.

Walt S. van der, Colbert S. C., Varoquaux G., 2011 The NumPy Array: A Structure for Efficient Numerical Computation. Comput. Sci. Eng. 13: 22–30.

